# Cross-Pipeline RNA-seq Analysis Reveals Core Regulatory Gene Signatures Driving P19 Cell Neurogenesis

**DOI:** 10.64898/2026.05.12.724245

**Authors:** Leila Rafiq, Hossein Khodadadi, Rim Drouzi, Mehdi Knidiri, Hiroaki Taniguchi

**Affiliations:** African Genome Center, University Mohammed VI Polytechnic (UM6P), Lot 660, Hay Moulay Rachid, 43150, Ben Guerir, Morocco; Institute of Genetics and Animal Biotechnology of the Polish Academy of Sciences, Postępu 36, Jastrzebiec 05-552, Poland

## Abstract

Understanding the mechanisms governing neuronal differentiation is essential for elucidating neurodevelopmental processes and identifying therapeutic targets for neurological disorders. In this study, we optimized serum-dependent induction conditions and benchmarked multiple RNA-seq pipelines to establish a robust in-vitro model of neurogenesis using P19 embryonal carcinoma cells. Retinoic acid (RA, 0.5 µM) was used to induce neuronal differentiation under varying concentrations (1%, 2%, and 5%) of fetal bovine serum (FBS) obtained from three suppliers. Morphological observation and marker gene analysis (MAP2, OCT4) revealed that serum concentration strongly influenced aggregation, survival, and neuronal commitment, with 2–5% FBS yielding optimal neurogenic differentiation. Total RNA extracted on day 10 of differentiation was subjected to RNA sequencing, and the resulting datasets were analyzed using four independent bioinformatics workflows: a Linux-based R pipeline (HISAT2 + featureCounts + DESeq2), nf-core, Galaxy, and BGI’s Dr. Tom platform. Differential gene expression analysis identified 9,943 differentially expressed genes (DEGs) (FDR < 0.05, |log2FC| > 1), enriched in synaptic assembly and axon development among upregulated genes, and in ribosome biogenesis and RNA processing among downregulated genes. Comparison across all pipelines revealed 62 consistently upregulated and 63 downregulated genes, representing a robust core signature of P19 neurogenesis. Together, these findings establish an optimized and reproducible framework for in-vitro neuronal differentiation and transcriptomic analysis, providing a foundation for mechanistic and disease-modeling studies in neurodevelopmental biology.

## II. INTRODUCTION

The nervous system is one of the most complex systems in the body, grouping a functional collection of cells, tissues, and organs that, together, regulate information processing and activity coordination in response to internal and external stimuli. In particular, the brain and spinal cord are part of the central nervous system (CNS), which is a highly organized and complex organ in mammals. This system is mainly composed of neurons and neuroglia which are highly specialized cells that transmit signals within the nervous system and between the latter and all parts of the body [1]. Hence, many diseases and brain disorders, such as Parkinson’s disease (PD), can be caused by a loss or dysfunction in these cells. Understanding the mechanisms of neurogenesis (i.e., how stem cells give rise to neurons) remains important for identifying regulatory pathways and potential therapeutic targets.

The most frequently used in-vitro models of embryonic neurogenesis are pluripotent stem cells (PSCs) [2], notably embryonic stem cells (ESCs) [3] and induced pluripotent stem cells (iPSCs) [4-6]. Their unlimited proliferative capacity as well as their ability to differentiate into the three germ layers (endoderm, mesoderm and ectoderm) make them very suitable for modeling early neurogenesis in vitro and for studying optimal conditions for their generation in laboratory [7]. However, these methods have issues due to their high cost and operational complexity.

P19 teratocarcinoma cells are known to possess the characteristic properties of embryonal carcinoma (EC) cells and can be maintained and propagated rapidly in tissue culture in an undifferentiated state [8]. P19 cells, much like embryonic stem cells, can differentiate into all three germ layers upon induction with different agents such as retinoic acid (RA) [8-10] and dimethyl sulfoxide (DMSO) [11], and also depending on the culture conditions [12]. Differentiation of these multipotential cells in vitro depends not only on the nature of the chemical inducer used or the culture condition, but also on the formation of cellular aggregates during the induction [8]. Exposure of aggregated cells to DMSO leads to their differentiation into endodermal and mesodermal derivatives such as cardiomyocytes and muscle cells [11]. While treatment with RA results in the formation of neuroectodermal cells including neuron-like cells, glia and fibroblast-like cells [11, 13].

Importantly, the amount of serum added to the cell culture media plays a key role in the cellular aggregation, thus affecting the neuronal differentiation [14, 15]. Fetal bovine serum (FBS) is commonly used as a low-cost supplement in cell-based applications, providing an optimal culture medium. FBS provides the cells with essential nutrients and growth factors that are necessary for maintaining them and helping with their growth [16]. It was introduced for the first time as a cellular growth stimulator, in the late 1950s, by Theodore Puck [17]. However, the use of FBS comes with some limitations such as the unknown exact composition or lot-to-lot variability even within the same company [18], which can heighten the risk of unexpected or undesired outcomes. Optimizing serum concentration is therefore necessary for reproducible differentiation and downstream molecular analyses.

The study of differential gene expression (DGE) is a main area of interest within the field of molecular biology. Understanding how cellular functions vary across different biological conditions is equivalent to comprehending how genes show different expression levels under varying conditions such as treated vs untreated, or wild-type vs mutated, etc. This can be achieved with the advent of next generation sequencing (NGS) and high throughput sequencing (HTS) technologies through transcriptome analysis. RNA sequencing (RNA-seq) has become an important tool for evaluating how cells modulate their gene expression, especially in response to neuronal induction. Understanding the molecular programs that govern neurogenesis requires analytical approaches capable of accurately detecting subtle changes in gene expression during this process. However, various widely used computational pipelines do exist (such as Linux command-line workflows, nf-core/rnaseq, Galaxy-based pipelines, and commercial or cloud-based solutions), yet the discrepancies offered by these approaches can yield divergent results even when applied to the same dataset, presenting a major challenge when studying neurogenesis. Therefore, a systematic comparison of these workflows in the context of neurogenesis is still needed.

In this study, we leveraged an optimized differentiation protocol, and we benchmarked multiple RNA-seq processing pipelines to assess how well each captures key transcriptomic transitions from pluripotency to neuronal fate and selected the most faithful workflow for identifying candidate genes that orchestrate neurogenesis in our model. This combined strategy provides a platform for dissecting neurodevelopmental mechanisms and for exploring disease-associated signatures in population-relevant contexts.

## III. MATERIALS AND METHODS

### Cell Culture and Differentiation Methods

#### Cell Culture Maintenance Condition

In a T25 cell culture plate (2 x 10^4^ cells/cm^2^), the P19 cells (donated by Katsuhiko Mikoshiba, RIKEN Center for Brain Science, Wako, Japan) were cultured with a maintenance medium containing DMEM High Glucose with Stable Glutamine and Sodium Pyruvate (Biowest, Catalogue number (CN): L0130) supplemented with 10% FBS, 100 units/mL penicillin and 100 units/mL streptomycin, and incubated at 37°C with 5% CO_2_.

#### Generation of Sphere Aggregates, Day 0-4 of Neurogenesis

Three flasks, each containing FBS from a different company, were used for culturing the P19 cells (1×10^6^ cells). Instead of the commercial names, they were mentioned as FBS A, FBS B, and FBS C. For the first 4 days, the cells (1×10^6^ cells) were plated on a 10 cm bacterial-grade Petri dish (Corning, Corning, NY, USA) with differentiation media (DMEM with 1, 2 and 5% FBS and 0.5 µM all-trans retinoic acid (Sigma, St. Louis, MO, USA)). After 2 days, sphere-shaped aggregates were observed. The differentiation medium was replaced with fresh medium, and the aggregates were seeded into a new 100 mm non-adherent culture dish and put back into the incubator for two more days. The embryoid-body forming cells were trypsinized and 2.5 x 10^6^ cells/well were seeded in the 6-well culture plate containing 3 mL of maintenance medium and incubated at 37°C with 5% CO_2_ until day 10 of neurogenesis.

#### Microscopic Observation

In order to track morphological changes, the formation of cell aggregates and neuronal cells was observed and pictured, respectively, at days 4 and 10 of RA-induced neurogenesis, via an inverted microscope (Nikon Eclipse TE2000-S) equipped with Nomarski differential interference contrast (DIC).

#### FBS Conditions and Suspension Culture Conditions

In order to validate RA-induced neurogenesis, three different FBS sources with three different dilutions (1%, 2%, and 5%) were used at the time of aggregate generation. Comparison of these conditions, regarding neurogenesis, was performed using RT-PCR for the stem marker OCT4 and the neuron marker MAP2.

### RNA Extraction, cDNA synthesis, RT-PCR, and RNA-seq

#### Isolation of RNA and PCR

On day 10 of neurogenesis, the total RNA was isolated from all Neuronal-like cells using the NucleoSpin RNA Mini kit for RNA purification (MACHEREY-NAGEL, CN: 740955.50). Next, 0.5 µg of total RNA was used to produce cDNA by the NG dART RT kit according to the manufacturer’s instructions (EURx, CN: E0801). And finally, PCR was performed for MAP2, OCT4, and β-Actin using RT HS-PCR Mix SYBR® kit (A&A Biotechnology, CN: 2017-100HS) under the following conditions: incubation at 98°C for 2 min, followed by 30 cycles of 98°C for 10s, 65°C for 15s, and 68°C for 1 min (Table 1).

**Table 1.**
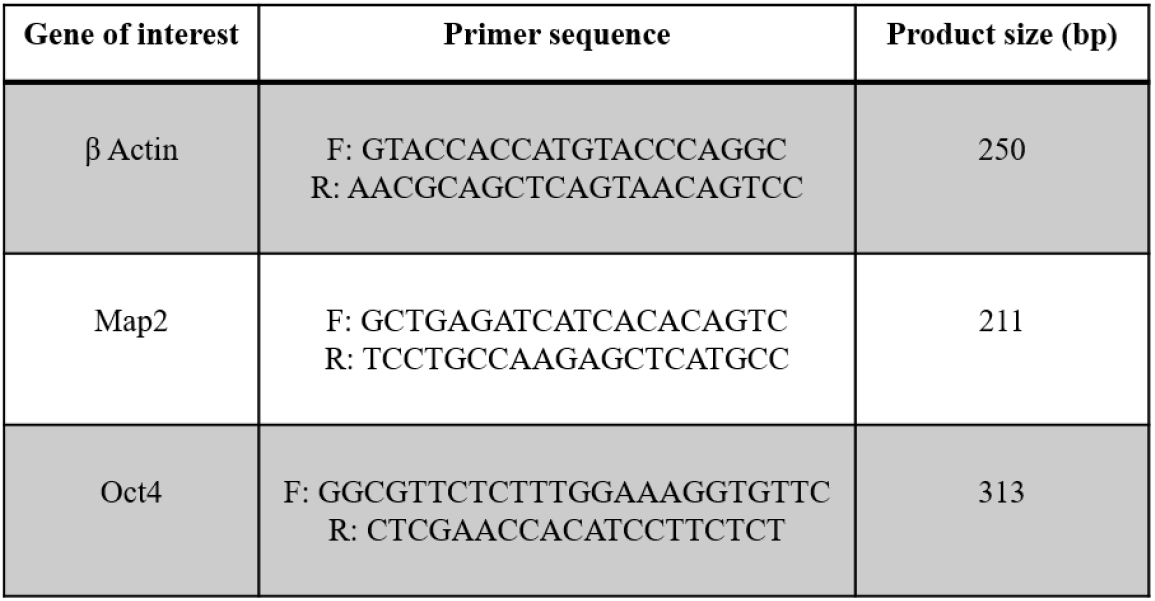
Primers used for PCR.

#### RNA-seq

Total RNA from neurons and WT groups was extracted and sent for high-throughput RNA sequencing performed by BGI (BGI Genomics, Shenzhen, China). Transcriptomics analysis, visualization, and interpretation were conducted using different pipelines.

#### Comparison of RNA-seq Analysis Pipelines

To evaluate performance, reproducibility and reliability, RNA-seq data was analyzed using four different pipelines, notably Linux- and R-based workflows, nf-core, Galaxy, and Dr. Tom, a web-based solution provided by BGI. As for Linux- and R-based workflows, we first performed quality control for the FASTQ raw reads in order to assess sequencing data quality, e.g., base quality, GC content, adaptor contamination, and duplicated reads. Afterward, the mouse reference genome (GRCm39) and corresponding GTF annotation were downloaded from GENCODE. Reads were aligned to the reference using HISAT2, a spliced alignment tool with efficient hierarchical indexing [19, 20], and the outputs were stored in a SAM (Sequence Alignment Map) format, then converted to BAM format using SAMtools [21, 22]. Gene-level read counts were then obtained with featureCounts, which summarizes mapped reads against annotated genomic features to produce a count matrix [22-24]. The latter is a raw count table which reports, for each sample, the number of sequencing reads that have been mapped to each gene [25, 26]. Rows contain ENSEMBL IDs [27] of expression features (genes or transcripts), and columns represent the sequenced samples. The values represent the number of reads mapped to each gene.

#### Differential Expression and Functional Analysis

The most important part of the present RNA-seq pipeline is differential gene expression (DGE) analysis. It is performed to infer a biological meaning to the analysis by determining which genes change their expression levels under different conditions [24]. This step is done after fragment (since we’re working with PE RNA-seq) counting and takes raw fragment counts contained in the expression matrix (from featureCounts) as input [28]. As the major goal of this analysis is to account for differences in gene expression level across the samples, it is necessary to normalize the counts before proceeding with the comparisons. Normalization adjusts for differences in gene length, read depth and technical biases, making comparison of gene expression between samples more accurate [29]. Differential gene expression (DGE) analysis was carried out in R using DESeq2, which models RNA-seq count data with a negative binomial distribution and applies the Median of Ratios normalization [30]. Enrichment analysis is a key step for understanding complex biological systems. This is a statistical analysis that looks for keywords of the set of molecules of interest in relation to a background set (reference genome or transcriptome) [31]. Functional enrichment was performed on significant DEGs (FDR < 0.05, |log2FC| > 1) using Gene Ontology (GO) terms covering biological process, molecular function, and cellular component [32, 33].

To benchmark results, we analyzed the same datasets with three additional workflows, using the same fastq files: and performed the following analysis. Nf-core/rnaseq (v3.18.0) is a curated Nextflow pipeline that integrates QC, alignment, quantification, and exploratory plots, ensuring reproducibility through containerization [34]. Galaxy (homepage: https://galaxyproject.org, main public server: https://usegalaxy.org), a user-friendly web-based platform where the workflow (FASTQC, HISAT2, DESeq2) was reproduced through a graphical interface [35]. Dr. Tom (BGI Genomics), an interactive platform providing DEG detection, enrichment, splicing, and literature-based annotation for genes of interest (https://www.bgi.com/global/dr-tom/).

## IV. RESULTS

### FBS Conditions and Cell Floating

After culture and treatment with RA of the P19 cells, on day 4 of the neurogenesis process, it was observed that in 5% FBS A, cells attached to the surface of the non-treated dish during the suspended culturing phase. In contrast, dilution of 1% FBS A and FBS C resulted in smaller aggregates formation in a reduced number. On the other hand, 1% of FBS B showed a comparable size and number of aggregates to those observed when using 2% FBS. For FBS B and C, 2 and 5% showed the same number and size of aggregates; however, in 5% FBS B, bigger aggregates formation was seen.

The number of cell survivals was evaluated on day 4 of the neurogenesis process, using an automated cell counter before seeding the cells on treated cell culture plates. It was found that in all FBS sources, the rate of alive cells was dramatically reduced in case of 1% FBS; however, in the 2% condition for FBS A and 5% condition for FBS B and FBS C, the highest cell viability was observed.

The cells were examined by light microscopy on day 10 of the neurogenesis process (Figure 1). It was observed that many cells died at 1% FBS for the 3 FBS sources, and only few neuron-like cells were formed. In case of 2% FBS, the formation of neuron-like cells was observed in fewer numbers compared to 5% FBS B and C (Figure 1).

**Figure 1.**
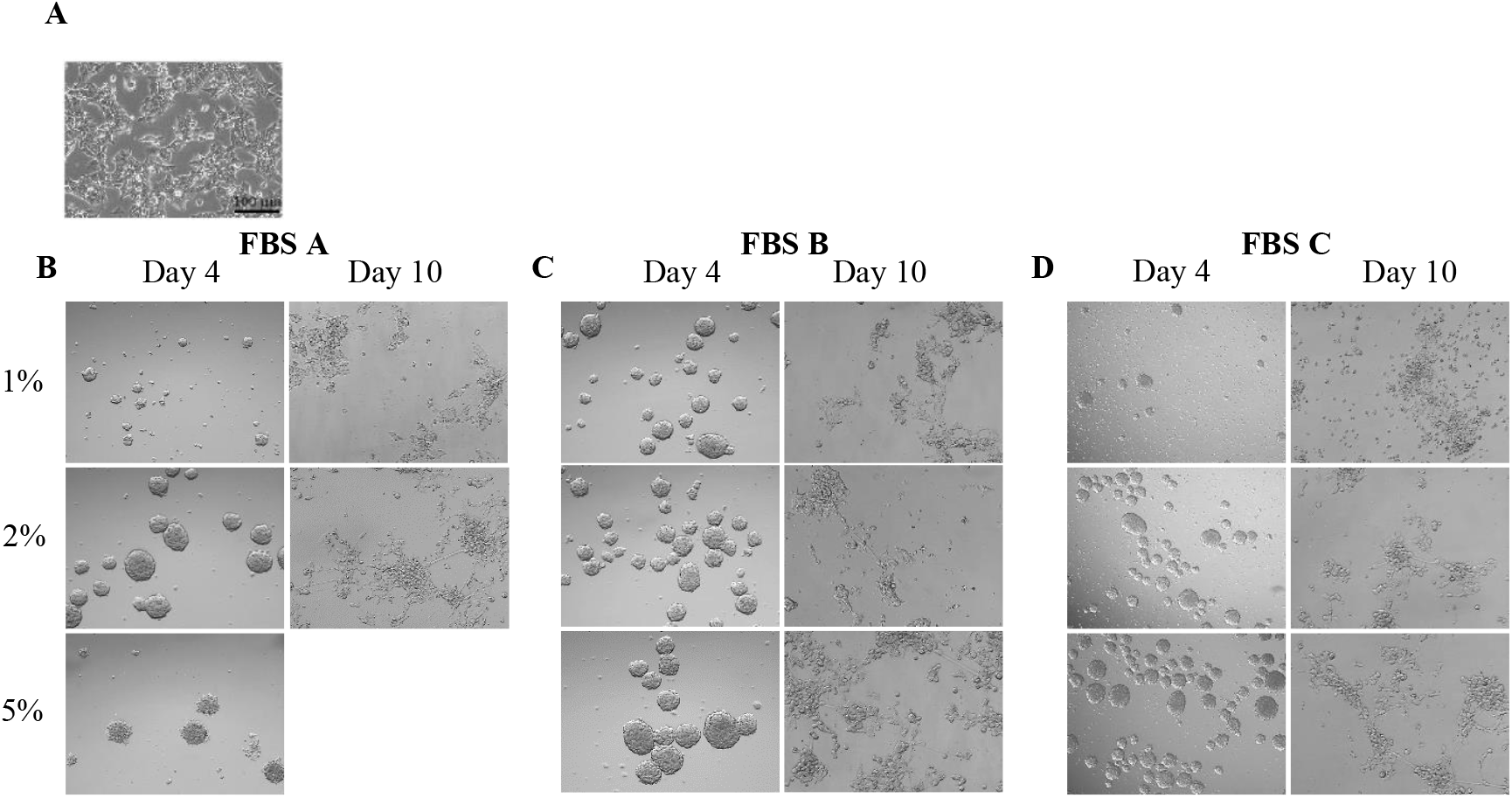
Representative images of analysis of P19 cell line RA-induced neurogenesis for three different FBS sources. **(A)** Light microscopic images of undifferentiated P19 cell line. **(B,C,D)** Light microscopic images of P19 cell line after 4 and 10 days of neurogenesis-following 4 days after RA-treatment (0.5µM) and cell aggregation stage with 1, 2, and 5% concentration of FBS. Scale bar = 100 µm

To validate these findings, the expression of MAP2 mRNA (a neuronal marker) was traced and compared to OCT4 (a pluripotent marker). For internal control, β-actin was used. On day 10 of neurogenesis, total RNA was extracted from all cells in different conditions of FBS in triplicate and was subjected to PCR. The results demonstrated a very weak trace of MAP2 expression in 1% FBS and a much higher expression in 2 and 5% FBS from the three companies. For further proof, the expression of OCT4 was evaluated by PCR, and no trace of this marker was observed at 2 and 5% FBS from the three sources. However, 1% FBS showed a faded signal, indicating the incomplete RA-induced neurogenesis (Figure 2).

**Figure 2.**
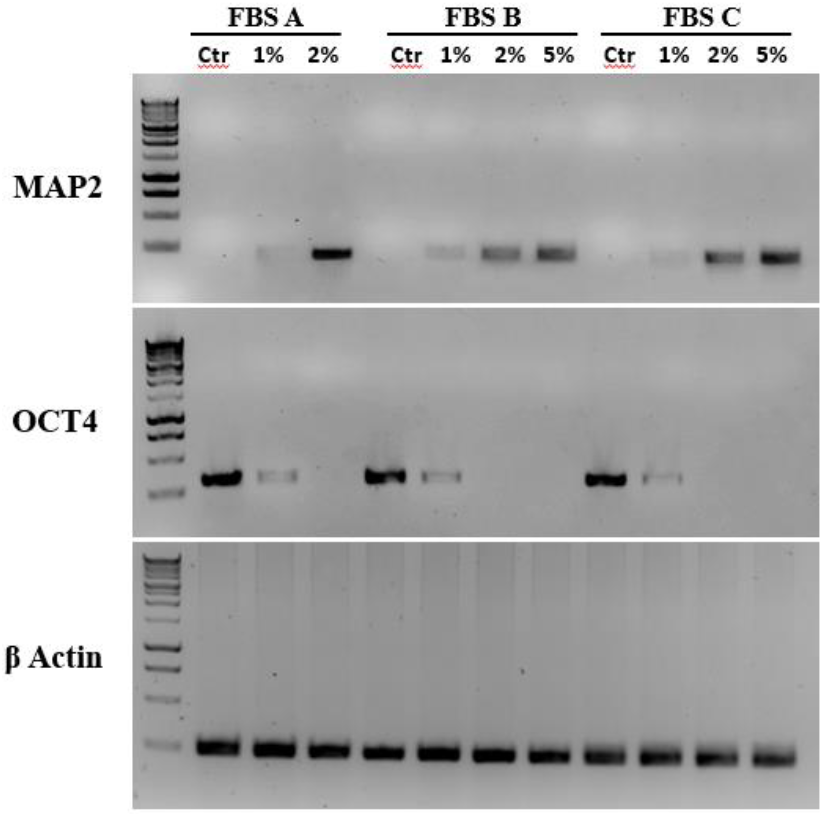
Changes of gene expression in P19 cell line. The band graph represents gene expression for undifferentiated P19 cell line as control (Ctr) for each FBS sources (FBS A, FBS B and FBS C) vs day 10 of RA-induced neurogenesis. *MAP2* gene expression was evaluated as a neuronal marker and OCT4 gene expression was evaluated as a pluripotent marker. β-Actin was used as the reference gene. Samples are loaded in the agarose gel (1.5%) in triple replications.

### RNA-seq Data Analysis

#### Identification of DEGs and Visualization

Differential gene expression (DGE) analysis is performed to infer a biological meaning to the analysis by determining which genes change their expression levels under different conditions [24]. This step is done after read (or fragment in case of PE RNA-seq) counting and takes raw read counts contained in the expression matrix as input [28]. The non-negative integer values in the matrix give an idea about how many fragments have been assigned to each gene in each sample. Thus, the RNA-seq experiment follows a discrete distribution of gene expression [36].

In order to account for differences in gene expression level across the samples, it is necessary to normalize the counts before proceeding with the comparisons. Normalization adjusts for differences in gene length, read depth and technical biases, making comparison of gene expression between samples more accurate [29].

In the present analysis, we use DESeq2 to detect the statistically significant DEGs between the two conditions using the NB model. The output from DESeq2 is formatted as a table of statistical measures for each gene, including log2FoldChange which is the magnitude of expression change between stem cells and neurons, pvalue, and padj which is the adjusted P-value or false discovery rate (FDR) for multiple testing correction (https://bioconductor.org/packages/release/bioc/vignettes/DESeq2/inst/doc/DESeq2.html). The results were filtered by setting two thresholds: padj < 0.05 and |log2FC| > 1, and a total of 9943 DEGs were identified, including 5791 upregulated and 4152 downregulated genes (|log2FC| > 1, FDR < 0.05).

##### GO Enrichment Analysis

Functional enrichment analysis is a method to explore biological functions of a set of genes and can be performed using a variety of tools, such as, Enrichr, or clusterProfiler. In our analysis, we performed GO functional enrichment analysis, using clusterProfiler as an R/Bioconductor package.

For the barplots in Figure 3 (A,C), the ontologies are color-coded to facilitate the comparison of the significantly enriched terms across them. Each GO term is represented by a bar for which the height is corresponding to the - log10(Adjusted P-value). This implies that more the length of the bar is high, the more the corresponding GO term is statistically significant (lower P-values). In addition, the GO terms are ordered in a way that the most significant ones appear at the top of the plot. In the dotplots Figure 3 (B,D), each enriched term is represented by a dot for which the size reflects the number of genes associated to that term, and the color varies depending on the Padj value.

**Figure 3.**
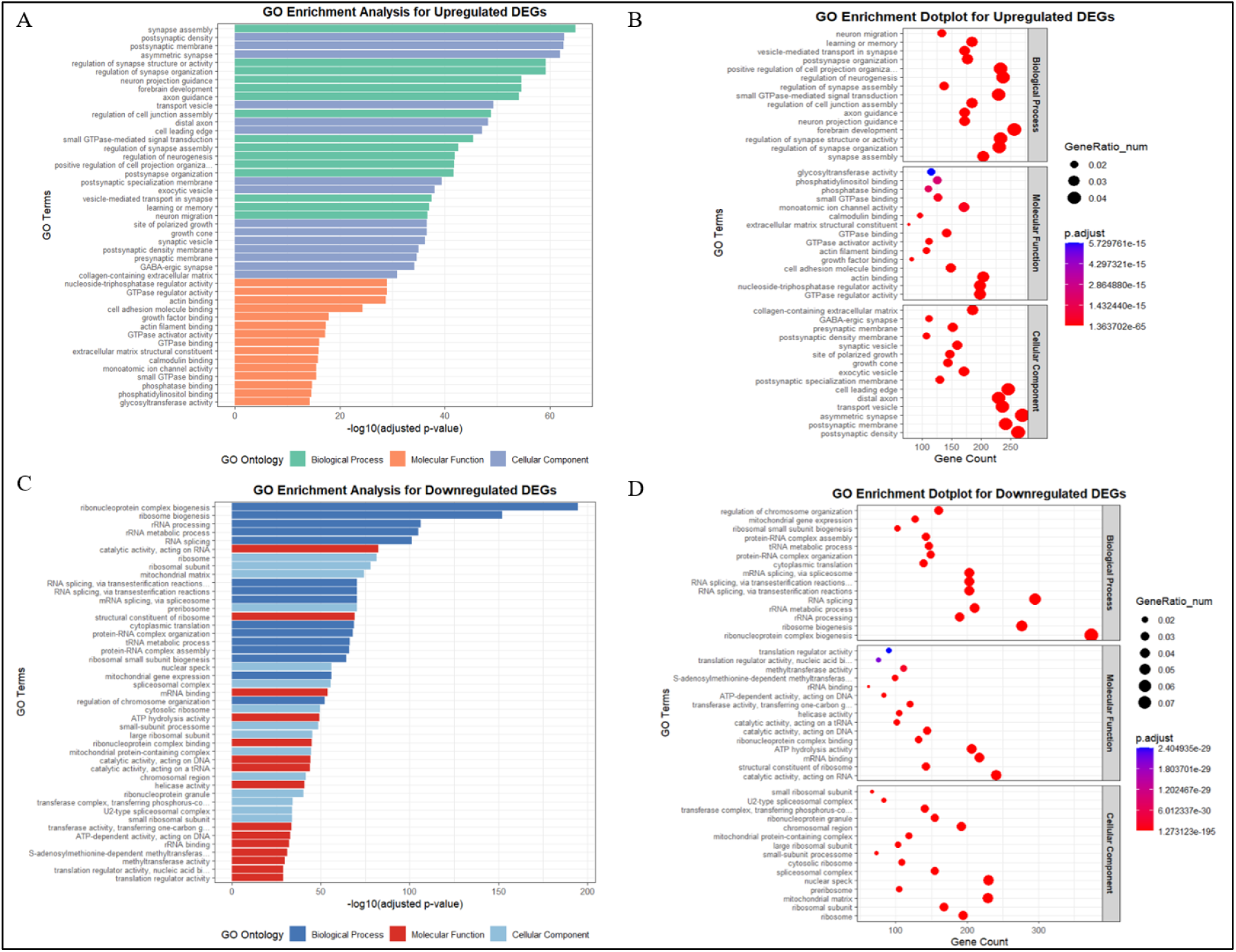
Integrated overview of GO results across three ontologies (BP, CC, and MF) for upregulated (A,B) and downregulated (C,D) DEGs. These plots present a visualization for the results obtained from GO enrichment analysis for upregulated and downregulated genes across three ontologies (BP, CC, and MF).

In the case of upregulated genes, the most significantly enriched BPs are synapse assembly, regulation of synapse structure or activity, and regulation of synapse organization suggesting a maturation of neurons and synapses. In the CC ontology, postsynaptic density, postsynaptic membrane, and asymmetric synapse are significantly enriched. Within the MF category, nucleoside triphosphatase regulator activity, GTPase regulator activity, as well was many functions related to biochemical binding are notably enriched (Figure 3A, B).

In the case of downregulated genes, among the BPs, the most enriched terms are ribonucleoprotein complex biogenesis, ribosome biogenesis, and rRNA processing. The most involved CCs are ribosome, ribosomal subunit, and mitochondrial matrix. In the MF ontology, terms such as catalytic activity acting on RNA, structural constituent of ribosome, and mRNA binding (Figure 3C, D).

#### Common Genes Obtained from the Four Methods

The first analysis using standard linux- and R-based workflows showed relevant biological insights. To confirm those results, the same analysis was performed using three other pipelines: nf-core, Galaxy, and Dr. Tom. A list of top100 DEGs, for both upregulated and downregulated genes separately, was generated from each pipeline, and Venn diagrams were created to compare the overlaps by showing the number of unique and shared genes that are consistently detected as differentially expressed by all the four pipelines.

The comparison revealed a core set of 62 common upregulated DEGs (Table 2, Figure 4) and 63 common downregulated ones (Table 3, Figure 5) consistently identified across all tested pipelines. Notably, Linux- and R-based analysis revealed the highest number of unique upregulated and downregulated genes, suggesting greater sensitivity. Galaxy and nf-core, on the other hand, showed the highest overlap with each other and with the other pipelines, especially Galaxy which detected very few numbers of unique genes, notably 3 upregulated and 13 downregulated unique DEGs (Figure 4, Figure 5).

**Table 2.** The Number of Overlapping Upregulated DEGs From the TOP100 Obtained by the Different Pipelines.

|  | BGI X<br>PIP1 | BGI X<br>NF-CORE | BGI X<br>GALAXY | PIP1 X<br>GALAXY | GALAXY X<br>NF-CORE | NF-CORE X<br>PIP1 | TOTAL<br>OVERLAP |
| --- | --- | --- | --- | --- | --- | --- | --- |
| Overlapping<br>Count | 66 | 86 | 82 | 79 | 83 | 67 | 62 |

**Table 3.** Overlapping Downregulated DEGs.

|  | BGI X<br>PIP1 | BGI X<br>NF-CORE | BGI X<br>GALAXY | PIP1 X<br>GALAXY | GALAXY X<br>NF-CORE | NF-CORE X<br>PIP1 | TOTAL<br>OVERLAP |
| --- | --- | --- | --- | --- | --- | --- | --- |
| Overlapping<br>Count | 67 | 76 | 71 | 81 | 74 | 73 | 63 |

**Figure 4.**
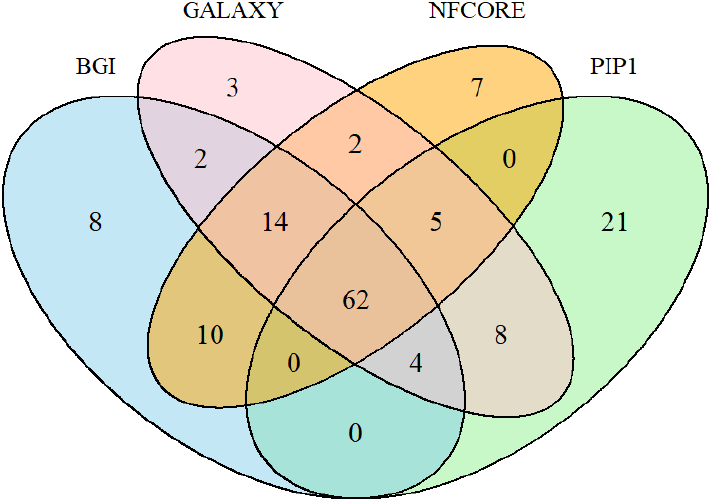
Venn Diagram Representing Intersections Between the 4 Pipelines, for the Top100 Upregulated DEGs. 62 is the total count for the overlapping genes between the 4 pipelines. In addition to the 62 genes that are common among the 4 pipelines, Galaxy found 3 other unique genes, while nf-core found 7, BGI Dr. Tom found 8, and the Linux-based pipeline found 21.

**Figure 5.**
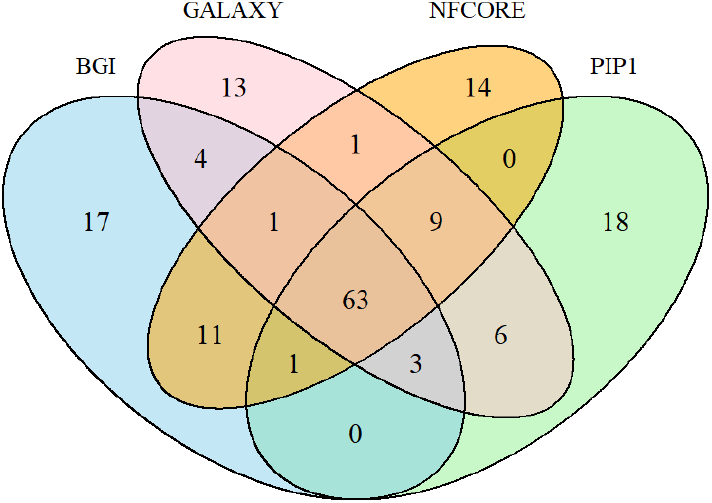
Venn Diagram Representing Intersections Between the 4 Pipelines, for the Top100 Downregulated DEGs. 63 is the total count for the overlapping genes between the 4 pipelines. In addition to the 63 genes that are common among the 4 pipelines, Galaxy found 13 other unique genes, while nf-core found 14, BGI Dr. Tom found 17, and the Linux-based pipeline found 18.

The intersection of the gene lists of the top 100 upregulated DEGs obtained from the 4 pipelines shows 62 overlapping genes in total (Table 4). As for the downregulated DEGs, 63 overlapping genes were extracted from the two tables of top 100 DEGs obtained from the four analyses (Table 5).

**Table 4.**
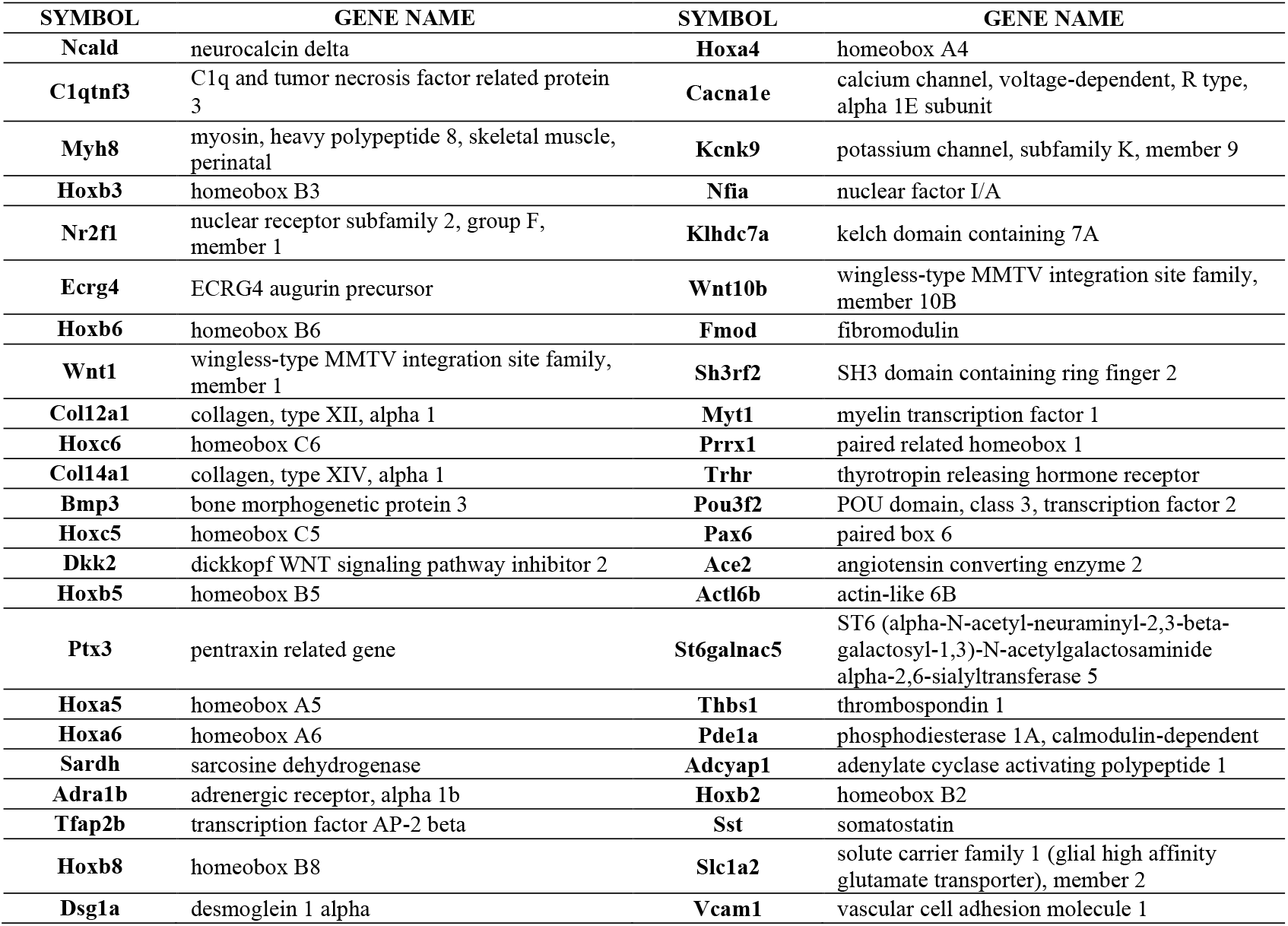
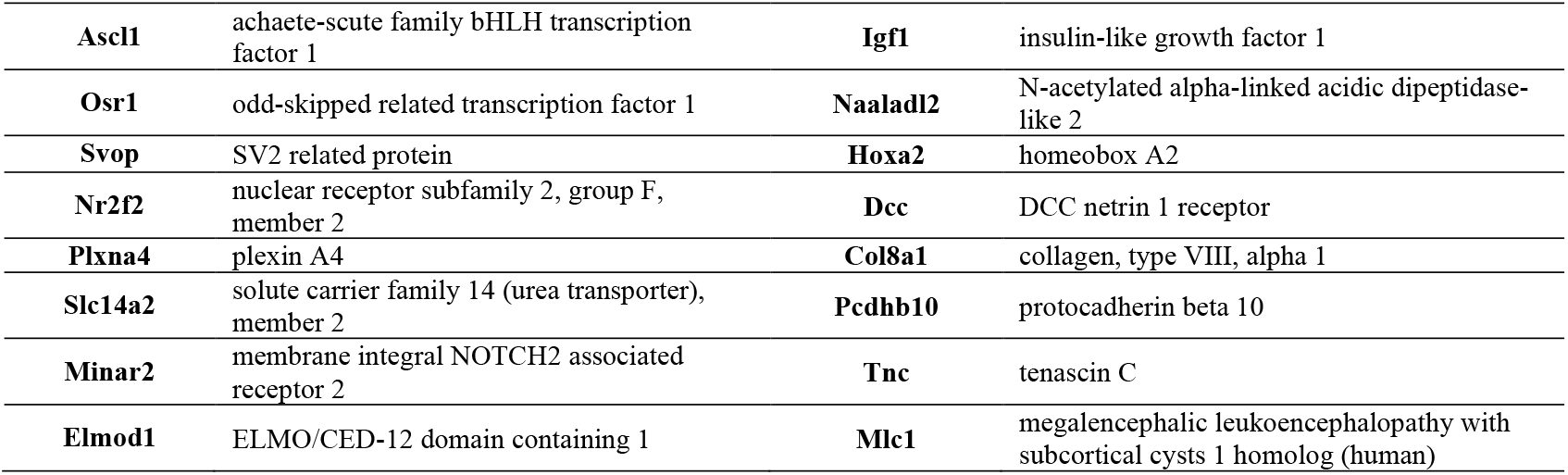
Overlapping Upregulated DEGs Annotated.

**Table 5.**
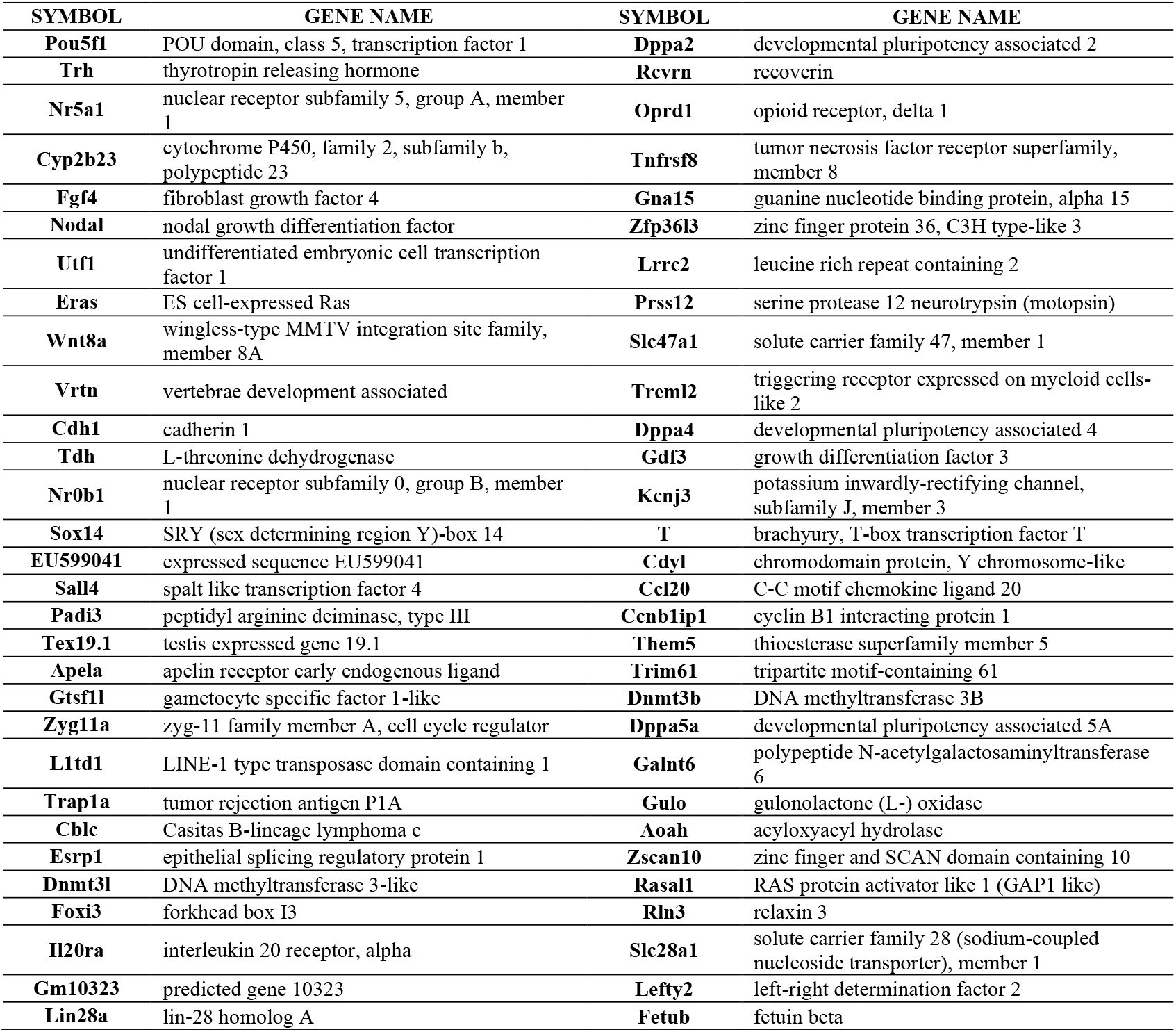
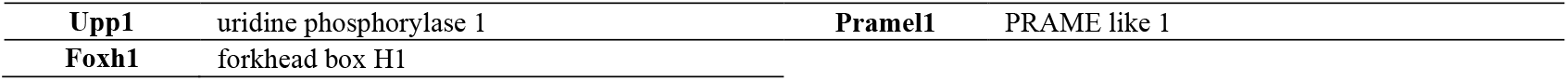
Overlapping Downregulated DEGs Annotated.

##### Overlapping DEGs

## V. DISCUSSION

This study provides an integrative view of the molecular events governing the transition from pluripotent to neuronal states in the P19 cell model. By combining optimized serum-dependent differentiation conditions with multi-pipeline RNA-seq benchmarking, we captured with high reproducibility the transcriptional “switch” that marks the loss of stem-cell identity and the activation of neuronal fate. Remarkably, four independent analytical workflows (Linux-based, nf-core, Galaxy, and BGI Dr. Tom) converged on an identical set of 62 up-regulated and 63 down-regulated genes, establishing one of the most consistent transcriptomic signatures observed in this P19 neurogenesis model. This reproducibility demonstrates that the observed gene-expression dynamics reflect a genuine biological program rather than computational artifacts.

A key achievement of this work is the establishment of a robust and convenient differentiation protocol that enables consistent neuronal induction. The results show that FBS source and dilution do influence aggregate size, viability, and neurogenesis efficiency. This suggests that the system is serum-tolerant but not serum-independent. While fetal bovine serum is known to introduce uncertainty due to compositional heterogeneity, our systematic comparison of three commercial FBS sources and multiple dilutions revealed that the combination of 0.5 µM retinoic acid with moderate serum levels (2–5%) reliably promotes neuronal differentiation and aggregate formation in P19 cells. This optimization mitigates one of the major practical limitations in neurogenesis studies which is dependence on specific serum lots and thus provides a standardized and reproducible approach for generating neuron-like cells in laboratories with limited access to defined media.

### 2% FBS is optimal for inducing neural differentiation of P19 cells

Upon induction by RA, many genes implicated in neurogenesis get activated. However, this process of cell differentiation into neurons is highly affected by aggregation formation, since this phenomenon allows for cell-cell interaction that is necessary in neurogenesis. Results have shown that different sources of FBS lead to different behaviors in cell cultures. Changing the source of FBS affects the size and the number of cell aggregates formed during RA-induced neurogenesis. Moreover, FBS dilution was demonstrated to be another important factor that plays a major role in aggregation formation.

Furthermore, it was also proved that FBS dilution affects cell survival. By evaluating the number of cell survivals, on day 4 of the neurogenesis process, the rate of alive cells varied dramatically when using different titrations of FBS, indicating that the dilution of FBS is crucial for cell survival during RA-induced cell aggregation.

On day 10 of the neurogenesis process, it was observed that FBS dilutions resulted in cell death in the case of 1% FBS, meanwhile neuron-like cells formation was observed in the case of 2% and 5% FBS, thus suggesting that 1% FBS is not recommended for RA-induced neurogenesis, while higher FBS dilutions, 2% and 5%, can lead to a complete formation of neuronal-like cells with high expression of neuronal markers and successful neurogenesis.

In the process of validating this hypothesis, the expression of MAP2 mRNA was traced and compared to that of OCT4. The results of PCR demonstrate that higher FBS dilutions, 2% and 5%, resulted in much higher expression of MAP2, compared to 1% which has shown a very weak trace of this neuronal marker. This finding aligned perfectly with the morphological observation of generated neurons. On the other hand, PCR evaluation of OCT4 expression showed that there was no trace of this marker in 2 and 5% FBS (from the 3 sources similarly), unlike 1% FBS, which indicates the incomplete RA-induced neurogenesis in the latter condition. Finally, 2% was chosen as the optimal condition, since it showed good morphological results with all FBS sources, and RNA was extracted from samples cultured in this condition and then sequenced.

The transcriptomic data strongly confirmed that 2% FBS is optimal for inducing neural differentiation of P19 cells. This induction led to the upregulation of many genes related to neurogenesis.

### Coordinated Transcriptional Switch from Pluripotency to Neuronal Fate

The most striking biological insight is the coordinated shutdown of pluripotency-maintenance machinery alongside the synchronous activation of neuronal-lineage regulators. Down-regulated genes such as Pou5f1 (OCT4), Sall4, Lin28a, Dppa4/5a, and Dnmt3b represent the molecular “off-switch” of stemness, indicating repression of self-renewal circuits and de-novo DNA-methylation activity. Their silencing suggests that RA-induced differentiation initiates epigenetic relaxation, allowing the activation of lineage-specific promoters. In contrast, up-regulated genes including Ascl1, Pax6, Pou3f2 (Brn2), Nfia, and Myt1 act as transcriptional “on-switches,” driving neuronal commitment and terminal differentiation. The concurrent induction of Wnt1, Wnt10b, Bmp3, and multiple Hox clusters (Hoxa2–a6, Hoxb2–b8, Hoxc5–c6) reveals the persistence of embryonic patterning cues that define spatial identity even in this simplified in-vitro system.

Beyond canonical regulators, several novel and rarely characterized genes such as Adcyap1 (PACAP) [37], Ncald (NCBI Gene: 83988), Minar2, and Ecrg4 were consistently up-regulated across all pipelines, suggesting previously unrecognized roles in neurogenesis. Adcyap1 (Adenylate cyclase activating polypeptide 1 or PACAP/Pituitary adenylate cyclase-activating polypeptide) is well studied in neuroscience, it encodes a neuropeptide with neuroprotective and synaptogenic effects, and it has been shown to promote the differentiation of embryonic neural stem cells into astrocytes, especially through the activation of the phospholipase C/protein kinase C (PKC) signaling pathway [37]; Ncald (or Neurocalcin Delta) and Minar2 (Major intrinsically disordered NOTCH2-binding receptor 1-like homolog) are implicated in calcium signaling and lipid metabolism [38, 39], and Ecrg4 (Esophageal cancer-related gene 4) encodes a protein called augurin that has been associated with neuronal survival and inflammation [40-42]. The reproducible enrichment of these genes implies that metabolic and secretory remodeling during neuronal differentiation involves broader signaling crosstalk than previously appreciated.

The global downregulation of ribosomal and mitochondrial genes further reflects a metabolic reprogramming from a proliferative to a quiescent oxidative state, an essential hallmark of neuronal maturation. The simultaneous repression of Dnmt3l/Dnmt3b (DNA methyltransferase enzymes) and Lin28a which encodes an RNA-binding protein that promotes pluripotency via regulation of miRNA biogenesis, with activation of the WNT–HOX–PAX6 transcriptional axis suggests a coordinated molecular hand-off between epigenetic silencing of stemness [43-48] and activation of morphogen-driven neuronal patterning [49-52]. This dual-switch mechanism represents a fundamental principle of lineage commitment: differentiation is not merely the gain of new gene expression but the active dismantling of the self-renewal network.

From a systems-biology perspective, the convergence of transcriptomic outputs across distinct computational platforms underscores the robustness of this model as a benchmark for neuronal differentiation studies. The reproducible identification of a conserved 125-gene core network provides a stable foundation for integrating future multi-omics layers, such as chromatin accessibility, proteomics, or single-cell transcriptomics, to dissect the temporal hierarchy of fate specification. Moreover, the combination of a serum-tolerant differentiation method with a cross-validated transcriptomic workflow establishes a practical and predictive platform for both mechanistic research and translational applications.

In summary, our findings reveal that RA-induced neurogenesis in P19 cells is governed by a synchronized molecular program in which the repression of stem-cell maintenance genes and activation of neuronal differentiation pathways occur in concert. The reproducibility and robustness of both the biological and computational frameworks highlight a predictive regulatory network linking DNMT3B–LIN28A silencing with WNT–HOX–PAX6 activation. Further investigation is needed to determine the functional contribution of these candidate regulators during neuronal lineage commitment and maturation.

Together with the establishment of an optimized serum-dependent differentiation protocol, this study offers both conceptual and methodological advances for modeling neurodevelopment, optimizing regenerative protocols, and exploring therapeutic gene networks underlying neuronal fate determination.

## VI. CONCLUSION

In this study, we demonstrated that P19 cells can be efficiently and reproducibly differentiated into neurons under 2% FBS in combination with retinoic acid. This condition supported robust aggregate formation, high survival, and clear acquisition of neuronal morphology. This differentiation at the molecular level was confirmed by the transcriptomic data analysis that showed strong upregulation of neurogenesis-associated genes and repression of pluripotency markers such as OCT4.

Comparison of four independent RNA-seq analysis pipelines revealed a highly consistent transcriptional signature with Galaxy and nf-core as the most reliable platforms with nf-core offering maximal reproducibility and automation for high-performance computing (HPC) environments, while Galaxy remains broadly accessible despite its longer processing time.

Together, these findings establish 2% FBS + RA as an optimized and serum-tolerant protocol for neuronal induction in P19 cells and provide a validated multi-pipeline RNA-seq framework for future neurogenesis studies. This dual biological and computational robustness contributes to a standardized platform that can support mechanistic research, multi-omics integration, and improved reproducibility in in-vitro neurodevelopmental modeling.

## Acknowledgement

The authors of this manuscript acknowledge that Gen AI has been employed in preparing this manuscript. Following established policies for using AI technology, the authors used ChatGPT, which belongs to Open AI for proofreading language in manuscripts. All suggestions from ChatGPT were reviewed and corrected by this group of authors in order to avoid inaccuracies in their manuscripts. This group of authors takes all responsibility for all information in this manuscript as well as all inaccuracies remaining.

## REFERENCES

1. Bazira, P.J., An overview of the nervous system. Surgery (Oxford), 2021. 39(8): p. 451–462.

2. Yap, M.S., et al., Neural Differentiation of Human Pluripotent Stem Cells for Nontherapeutic Applications: Toxicology, Pharmacology, and In Vitro Disease Modeling. Stem Cells Int, 2015. 2015: p. 105172.

3. Zhang, S.C., et al., In vitro differentiation of transplantable neural precursors from human embryonic stem cells. Nat Biotechnol, 2001. 19(12): p. 1129–33.

4. Lu, J., et al., Application of epigenome-modifying small molecules in induced pluripotent stem cells. Med Res Rev, 2013. 33(4): p. 790–822.

5. Masotti, A., et al., Aged iPSCs display an uncommon mitochondrial appearance and fail to undergo in vitro neurogenesis. Aging (Albany NY), 2014. 6(12): p. 1094–108.

6. Velasco, I., et al., Concise review: Generation of neurons from somatic cells of healthy individuals and neurological patients through induced pluripotency or direct conversion. Stem Cells, 2014. 32(11): p. 2811–7.

7. Azari, H. and B.A. Reynolds, In Vitro Models for Neurogenesis. Cold Spring Harb Perspect Biol, 2016. 8(6).

8. Bain, G., et al., From embryonal carcinoma cells to neurons: the P19 pathway. Bioessays, 1994. 16(5): p. 343–8.

9. Suzuki, Y., et al., Identification of differentially expressed mRNAs during neuronal differentiation of P19 embryonal carcinoma cells. Neuroscience Research, 1995. 23(1): p. 65–71.

10. Yao, M., G. Bain, and D.I. Gottlieb, Neuronal differentiation of P19 embryonal carcinoma cells in defined media. J Neurosci Res, 1995. 41(6): p. 792–804.

11. McBurney, M.W., et al., Control of muscle and neuronal differentiation in a cultured embryonal carcinoma cell line. Nature, 1982. 299(5879): p. 165–167.

12. Wei, Y., T. Harris, and G. Childs, Global gene expression patterns during neural differentiation of P19 embryonic carcinoma cells. Differentiation, 2002. 70(4-5): p. 204–19.

13. Jones-Villeneuve, E.M., et al., Retinoic acid induces embryonal carcinoma cells to differentiate into neurons and glial cells. J Cell Biol, 1982. 94(2): p. 253–62.

14. Liu, Q., et al., [Effect of serum on the differentiation of neural stem cells]. Zhongguo Xiu Fu Chong Jian Wai Ke Za Zhi, 2018. 32(2): p. 223–227.

15. Khasawneh, R.R., et al., Addressing the impact of different fetal bovine serum percentages on mesenchymal stem cells biological performance. Mol Biol Rep, 2019. 46(4): p. 4437–4441.

16. Petrović, D.J., et al., Effect of Fetal Bovine Serum or Basic Fibroblast Growth Factor on Cell Survival and the Proliferation of Neural Stem Cells: The Influence of Homocysteine Treatment. Int J Mol Sci, 2023. 24(18).

17. Puck, T.T., S.J. Cieciura, and A. Robinson, Genetics of somatic mammalian cells. III. Long-term cultivation of euploid cells from human and animal subjects. J Exp Med, 1958. 108(6): p. 945–56.

18. Baker, M., 1,500 scientists lift the lid on reproducibility. Nature, 2016. 533(7604): p. 452–454.

19. Kim, D., B. Langmead, and S.L. Salzberg, HISAT: a fast spliced aligner with low memory requirements. Nat Methods, 2015. 12(4): p. 357–60.

20. Kim, D., et al., Graph-based genome alignment and genotyping with HISAT2 and HISAT-genotype. Nat Biotechnol, 2019. 37(8): p. 907–915.

21. Li, H., et al., The Sequence Alignment/Map format and SAMtools. Bioinformatics, 2009. 25(16): p. 2078–9.

22. Liao, Y., G.K. Smyth, and W. Shi, featureCounts: an efficient general purpose program for assigning sequence reads to genomic features. Bioinformatics, 2014. 30(7): p. 923–30.

23. Conesa, A., et al., A survey of best practices for RNA-seq data analysis. Genome Biol, 2016. 17: p. 13.

24. Stark, R., M. Grzelak, and J. Hadfield, RNA sequencing: the teenage years. Nature Reviews Genetics, 2019. 20(11): p. 631–656.

25. Love, M., et al., RNA-Seq workflow: gene-level exploratory analysis and differential expression [version 1;peer review: 2 approved]. 2015. 4(1070).

26. Koch, C.M., et al., A Beginner’s Guide to Analysis of RNA Sequencing Data. Am J Respir Cell Mol Biol, 2018. 59(2): p. 145–157.

27. Flicek, P., et al., Ensembl 2012. Nucleic Acids Res, 2012. 40(Database issue): p. D84–90.

28. Oshlack, A., M.D. Robinson, and M.D. Young, From RNA-seq reads to differential expression results. Genome Biology, 2010. 11(12): p. 220.

29. Kvam, V.M., P. Liu, and Y. Si, A comparison of statistical methods for detecting differentially expressed genes from RNA-seq data. Am J Bot, 2012. 99(2): p. 248–56.

30. Love, M.I., W. Huber, and S. Anders, Moderated estimation of fold change and dispersion for RNA-seq data with DESeq2. Genome Biol, 2014. 15(12): p. 550.

31. Marco-Ramell, A., et al., Evaluation and comparison of bioinformatic tools for the enrichment analysis of metabolomics data. BMC Bioinformatics, 2018. 19(1): p. 1.

32. The Gene Ontology Resource: 20 years and still GOing strong. Nucleic Acids Res, 2019. 47(D1): p. D330–d338.

33. Ashburner, M., et al., Gene ontology: tool for the unification of biology. The Gene Ontology Consortium. Nat Genet, 2000. 25(1): p. 25–9.

34. Langer, B.E., et al., Empowering bioinformatics communities with Nextflow and nf-core. Genome Biology, 2025. 26(1): p. 228.

35. Afgan, E., et al., The Galaxy platform for accessible, reproducible and collaborative biomedical analyses: 2018 update. Nucleic Acids Res, 2018. 46(W1): p. W537–w544.

36. Soneson, C. and M. Delorenzi, A comparison of methods for differential expression analysis of RNA-seq data. BMC Bioinformatics, 2013. 14: p. 91.

37. Watanabe, J., et al., Pituitary adenylate cyclase-activating polypeptide-induced differentiation of embryonic neural stem cells into astrocytes is mediated via the beta isoform of protein kinase C. J Neurosci Res, 2006. 84(8): p. 1645–55.

38. Upadhyay, A., et al., Neurocalcin Delta Knockout Impairs Adult Neurogenesis Whereas Half Reduction Is Not Pathological. Front Mol Neurosci, 2019. 12: p. 19.

39. Gao, G., et al., Kiaa1024L/Minar2 is essential for hearing by regulating cholesterol distribution in hair bundles. Elife, 2022. 11.

40. Gonzalez, A.M., et al., Ecrg4 expression and its product augurin in the choroid plexus: impact on fetal brain development, cerebrospinal fluid homeostasis and neuroprogenitor cell response to CNS injury. Fluids Barriers CNS, 2011. 8(1): p. 6.

41. Nakatani, Y., H. Kiyonari, and T. Kondo, Ecrg4 deficiency extends the replicative capacity of neural stem cells in a Foxg1-dependent manner. Development, 2019. 146(4).

42. Yang, K., et al., Downregulation of ECRG4 by DNMT1 promotes EC growth via IRF3/IFN-γ/miR-29b/DNMT1/ECRG4 positive feedback loop. iScience, 2025. 28(1): p. 111614.

43. Bourc’his, D., et al., Dnmt3L and the establishment of maternal genomic imprints. Science, 2001. 294(5551): p. 2536–9.

44. Bourc’his, D. and T.H. Bestor, Meiotic catastrophe and retrotransposon reactivation in male germ cells lacking Dnmt3L. Nature, 2004. 431(7004): p. 96–9.

45. Neri, F., et al., Dnmt3L antagonizes DNA methylation at bivalent promoters and favors DNA methylation at gene bodies in ESCs. Cell, 2013. 155(1): p. 121–34.

46. Okano, M., et al., DNA methyltransferases Dnmt3a and Dnmt3b are essential for de novo methylation and mammalian development. Cell, 1999. 99(3): p. 247–57.

47. Tsialikas, J. and J. Romer-Seibert, LIN28: roles and regulation in development and beyond. Development, 2015. 142(14): p. 2397–404.

48. Zhang, J., et al., LIN28 Regulates Stem Cell Metabolism and Conversion to Primed Pluripotency. Cell Stem Cell, 2016. 19(1): p. 66–80.

49. Lekven, A.C., et al., Zebrafish wnt8 encodes two wnt8 proteins on a bicistronic transcript and is required for mesoderm and neurectoderm patterning. Dev Cell, 2001. 1(1): p. 103–14.

50. Mallo, M., D.M. Wellik, and J. Deschamps, Hox genes and regional patterning of the vertebrate body plan. Dev Biol, 2010. 344(1): p. 7–15.

51. St-Onge, L., et al., Pax6 is required for differentiation of glucagon-producing alpha-cells in mouse pancreas. Nature, 1997. 387(6631): p. 406–9.

52. Götz, M., A. Stoykova, and P. Gruss, Pax6 controls radial glia differentiation in the cerebral cortex. Neuron, 1998. 21(5): p. 1031–44.

